# Demographic responses of oceanic island birds to local and regional ecological disruptions revealed by whole-genome sequencing

**DOI:** 10.1101/2023.09.12.555930

**Authors:** Maëva Gabrielli, Thibault Leroy, Jordi Salmona, Benoit Nabholz, Borja Milá, Christophe Thébaud

## Abstract

Disentangling the effects of ecological disruptions operating at different spatial and temporal scales in shaping past species’ demography is particularly important in the current context of rapid environmental changes driven by both local and regional factors. We argue that volcanic oceanic islands provide useful settings to study the influence of past ecological disruptions operating at local and regional scales on population demographic histories. We investigate potential drivers of past population dynamics for three closely related species of passerine birds from two volcanic oceanic islands, Reunion and Mauritius (Mascarene archipelago), with distinct volcanic history. Using ABC and PSMC inferences from complete genomes, we reconstructed the demographic history of the Reunion Grey White-eye (*Zosterops borbonicus* (Pennant, 1781)), the Reunion Olive White-eye (*Z. olivaceus* (Linnaeus, 1766)), and the Mauritius Grey White-eye (*Z. mauritianus* (Gmelin, 1789)), and searched for possible causes underlying similarities or differences between species living on the same or different islands. Both demographic inferences strongly support ancient and long-term expansions in all species. They also reveal different trajectories between species inhabiting different islands, but consistent demographic trajectories in species or populations from the same island. Species from Reunion appear to have experienced synchronous reductions in population size during the Last Glacial Maximum, a trend not seen in the Mauritian species. Overall, this study suggests that local events may have played a role in shaping population trajectories of these island species. It also highlights the potential of our conceptual framework to disentangle the effects of local and regional drivers on past species’ demography and long-term population processes.

## Introduction

Ecological disruptions due to variations in environmental conditions can have profound effects on local population size by directly affecting individual physiology or indirectly modifying population niche limits. These disruptions can be triggered by major environmental events, such as global climatic changes, that can lead to drastic reductions in population size. Such bottlenecks have important consequences for gene pools owing to their effect on effective population sizes (*N_e_*) (Maruyama and Fuerst 1985; Nei, Maruyama, and Chakraborty 1975; Tajima 1989; Watterson 1984). Such effects have been documented across different taxa in the case of past global climatic events, implying some synchrony in population fluctuations at a very large spatial scale in response to climatic changes (Chattopadhyay et al. 2019; Germain et al. 2023). In turn, severe ecological disruptions at the local level can have more temporary and localized effects on populations. These include volcanic eruptions, forest fires, and direct or indirect human impacts on the environment through predation or habitat modification, all of which have the capacity to radically transform local landscapes (Dornelas 2010). Distinguishing the effects of local-scale drivers relative to large-scale processes in shaping the demography of populations and species over time, especially when both occurred in the past before human impact, might be essential if we want to predict accurately how human-driven climate change will challenge the global persistence of many species and their conservation (Nogués-Bravo et al. 2018; Urban et al. 2016). Yet, evaluating the relative importance of local-based and global-based influences remains challenging since past local effects are generally poorly understood at the scale of a species’ range and can be confounded by global effects.

Volcanic oceanic islands possess a number of features that could help to solve problems associated with the difficulty of distinguishing the effects of past ecological disruptions operating at local versus regional scales on population demographic histories (Warren et al. 2015). First, they have usually undergone a series of localized processes, such as major geodynamic events (Whittaker, Triantis, and Ladle 2008) or human impact, but have also experienced the effects of past changes in the global climate (Weigelt and Kreft 2013). Second, most volcanic oceanic islands are configured into groups of islands that are closely spaced and have similar origins, geophysical settings and climates. Yet, each island within such volcanic archipelagos has its own geological life cycle (Whittaker et al. 2008), and different islands have experienced independent bouts of ecological disruptions, mainly owing to volcanic episodes and related geodynamic processes that are often asynchronous across islands. Therefore, studying volcanic archipelagos with islands at different stages of their life cycle could allow for assessing the relative importance of local (*i.e.*, island-specific) and regional (*i.e.*, at the archipelago scale) disruption events on population size trajectories by comparing long-term demographic responses of species living on the same or different islands within an archipelago (Figure 1). In addition, the isolated nature of these islands is an asset for inferring demographic histories of insular lineages as most species are island endemics and interference from external source populations that can generate complex demographic scenarios, as is often found in continental settings, is expected to be minimal (Losos and Ricklefs 2009). Furthermore, because different islands often host allopatric sister species, archipelagos allow for the comparison of demographic histories from immediately related species, minimizing possible higher-level confounding effects (Weir and Lawson 2015).

**Figure 1:**
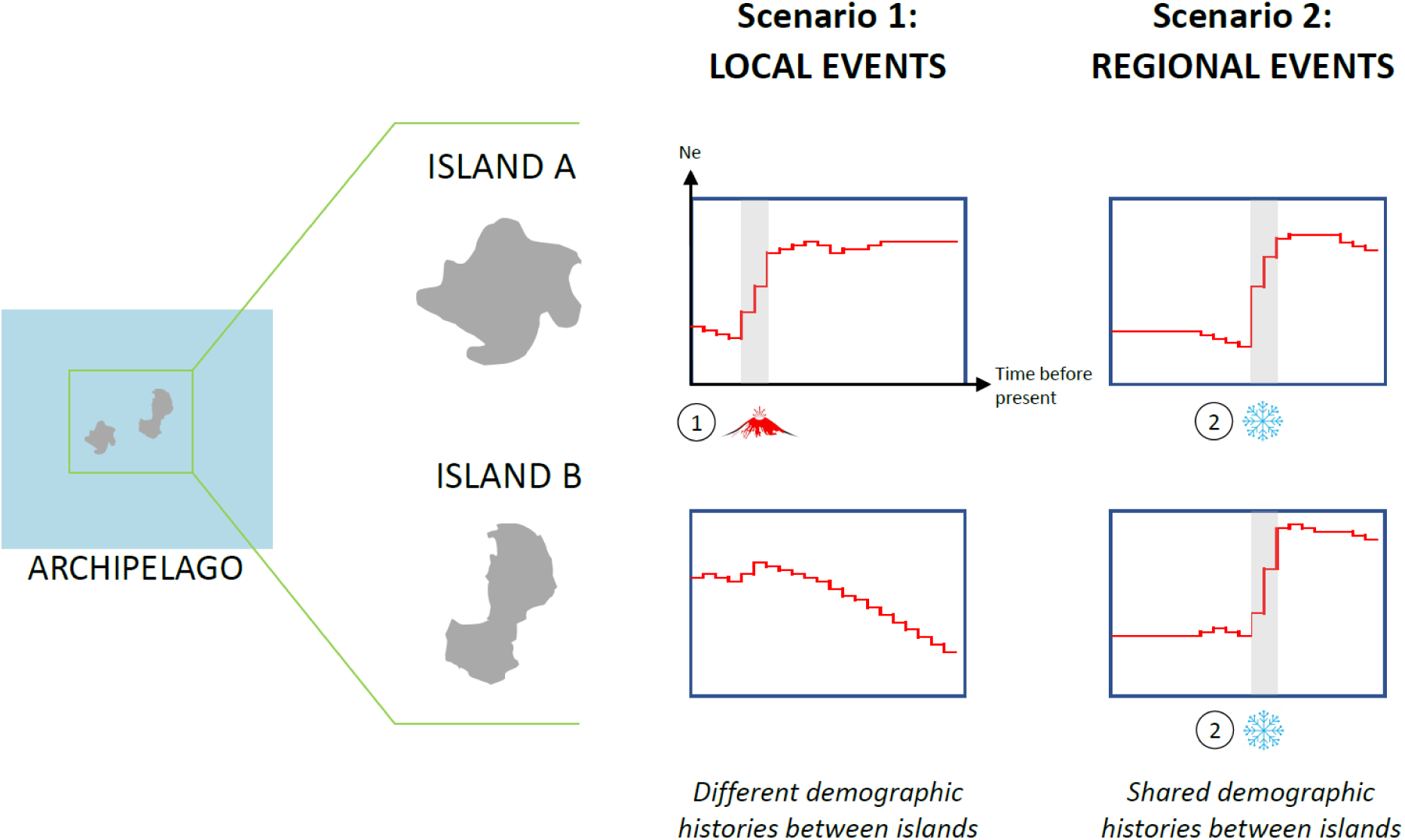
Proposed strategy to compare the effects of local events and regional events on genetic diversity by comparing the demographic histories of closely related populations or species inhabiting different islands from the same volcanic oceanic archipelago. Plots represent effective population size (*N_e_*) through time for a population or species, and numbers indicate different events of ecological disruption associated with bottlenecks: event 1 is a volcanic eruption on island A while event 2 corresponds to a regional drop in temperatures, as observed during a glacial maximum, that affects demographic histories on islands A and B synchronously, all else being equal. Contemporary bottlenecks suggest the importance of events at a large geographic scale while bottlenecks associated with island-specific events suggest the importance of local events as drivers of population histories.

Some studies have inferred the demographic histories of species from volcanic archipelagos (*e.g.* (Lamichhaney et al. 2015; Martin et al. 2021), but did not attempt to compare the population histories of species living on the same and different islands. Here, we propose to do so using, as a study system, the Mascarene archipelago, a remote island group in the southwestern Indian Ocean composed of two main islands, Mauritius and Reunion. The two islands are separated by 140 km of deep water and were never connected to each other (Thébaud et al. 2009). They are comparable in size while being at different stages of island ontogeny (see Fernández-Palacios et al., 2011), and therefore exhibit very different topographies: Reunion, a younger island (2 my) is in the erosion and landslide phase and has a very rugged landscape, reaching an elevation of 3,071 m, while Mauritius, an older island (8 my) in the late basal plain phase, has experienced the long-term effects of erosion and culminates at just 828 m (Duncan 2009). Global climatic events, including Late Quaternary events, appear to have affected species’ demographic histories on both islands (de Boer et al. 2015; Garot et al. 2019; Heller et al. 2008; Salmona et al. 2012; Salmona, Heller, Quéméré, et al. 2017). However, the intensity of ecological disruption as experienced by animal and plant populations may have been very different on the two islands since the persistence of possible refugia during extreme cooling events is positively related to mountainous topography (Harter et al., 2015; Mastretta-Yanes et al., 2015; but see Salces-Castellano et al., 2021). Localized and temporary, island-specific events likely to have affected many species’ population trajectories have also occurred on both islands (Cheke and Hume 2010). In particular a series of large explosive volcanic eruptions that took place around 200,000 ya (198,800 ya ± 2,500 years) on Reunion (Castellanos Melendez et al. 2023), and a series of basaltic lava flows that have occurred regularly (on average every 21,000 years) in the last 500,000 years on Mauritius, which currently cover nearly 75% of the island (Moore et al. 2011).

To illustrate potential effects of local and regional disruption events on population demographic trajectories, we focused on three species of white-eyes (Zosteropidae), all single-island endemics in the Mascarene Archipelago (Milá et al., 2010; Warren et al., 2006; Figure 2). These small passerine birds show limited dispersal capacity (Bertrand et al., 2014; Chris Smith, pers. comm.), a feature that could make them sensitive to major climatic and environmental changes (Germain et al. 2023). We hypothesized that (1) if island-specific events had major effects on species’ demography, we should recover island-specific demographic signals shared by species living on a particular island, and similarly, (2) if regional climatic events had severe impacts on species at the scale of the entire archipelago, we should find similar signals of demographic events between species living on different islands (Figure 1). Taking advantage of the recent public release of a high-quality reference genome for the Reunion Grey White-eye (*Zosterops borbonicus* (Pennant, 1781)) (Leroy, Anselmetti, et al. 2021), we used shotgun sequencing to generate whole-genome sequence data from the Reunion Grey White-eye, the Reunion Olive White-eye (*Zosterops olivaceus* (Linnaeus, 1766)) and the Mauritius Grey White-eye (*Zosterops mauritianus* (Gmelin, 1789)). The first two species are endemic to Reunion whereas the third species is endemic to Mauritius. All three species are immediate relatives (Milá et al., 2010; Warren et al., 2006; Figure 2) and are common and widely distributed in their respective islands. Therefore, it seems reasonable to assume that their long-term population dynamics have been little affected by anthropogenic activities since humans colonized these islands (Cheke and Hume 2010). In addition, the Reunion Grey White-eye is represented by multiple geographic forms that originated and diversified within Reunion (Gabrielli et al. 2020; Gill 1973). Three forms are restricted to the lowlands, whereas a fourth form is only found in the highlands, primarily between 1,400 and 3,000 m (Bertrand et al. 2016; Cornuault et al. 2015). These forms can therefore be considered as replicates of lineages from the same island but also as indicators of within-island environmental differences. To reconstruct the long-term trajectory of effective population sizes (Emerson, Paradis, and Thébaud 2001), we first used a series of Markovian coalescent analyses at the individual level. Although these approaches have been intensively used in the past decade (Li and Durbin 2011; Schiffels and Durbin 2014; Terhorst, Kamm, and Song 2017), a potentially serious limitation to using Markovian coalescent methods is that they assume that the focal species behaves as a panmictic unit (Mazet et al. 2016). Therefore, we also used demographic inferences based on Approximate Bayesian Computation (ABC; Beaumont et al., 2002), since they allow the evaluation of various scenarios of population size changes accounting for periods of gene exchange and estimate the timing of demographic events. Our results strongly support ancient expansions in the three species, likely associated with evolutionary processes that took place on the islands themselves since they diverged from an ancestral stock. Furthermore, our results indicate that the two species of Reunion white-eyes similar demographic histories, distinct from the demographic history of the Mauritius Grey White-eye. These findings suggest that local events have played a rather significant role in shaping population trajectories in Mascarene white-eyes. Our conceptual framework and its working hypotheses appear to provide an improved basis for disentangling the effects of local and regional drivers on past demography of species and a better understanding of population processes that are of paramount conservation importance in the current context of rapid environmental change driven by both local and regional factors of ecological disruption.

**Figure 2:**
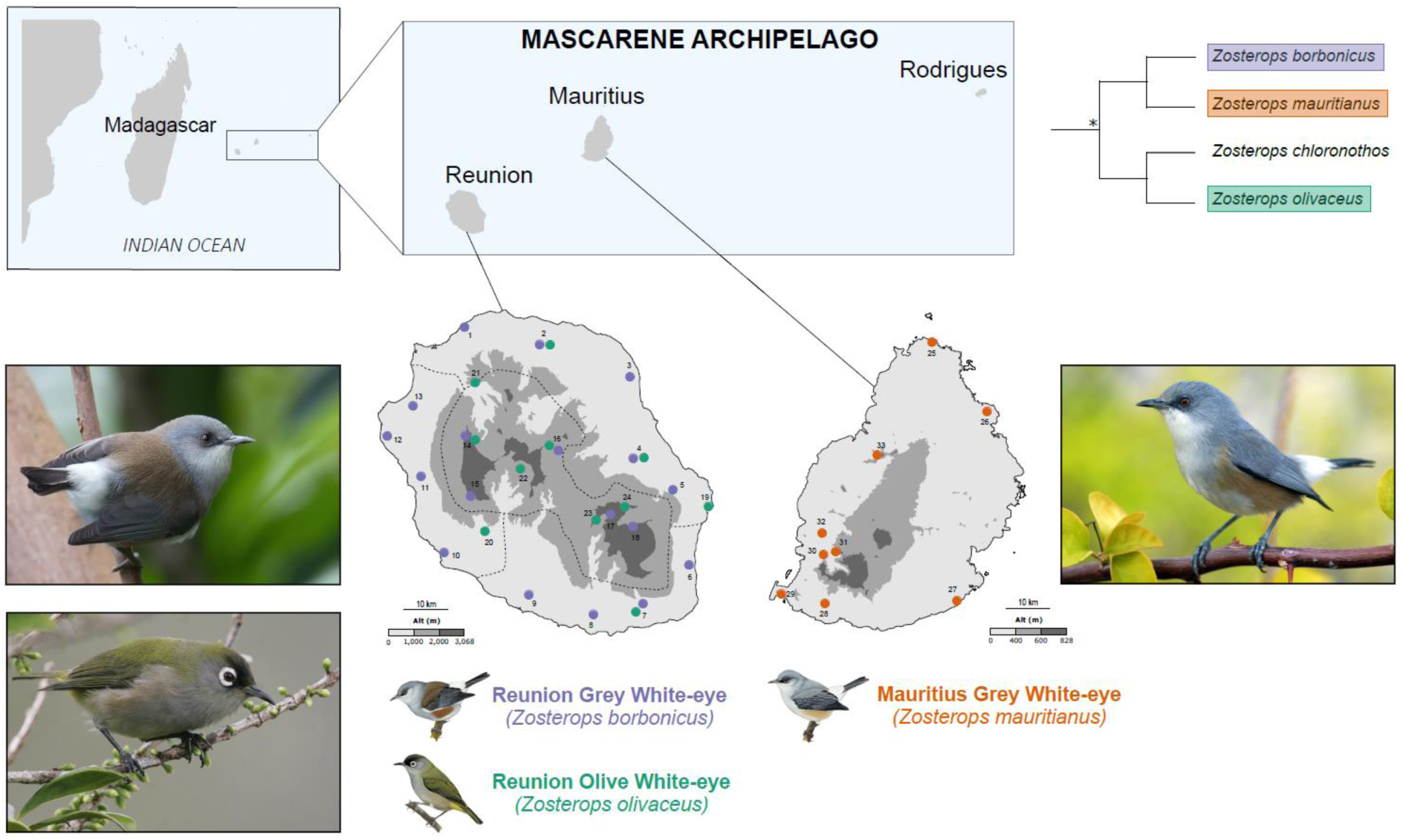
Distribution range and sampling localities. Localities are labelled with numbers (see Table S1 for names and coordinates) and circles with different colours represent samples from Reunion Grey White-eye (purple), Reunion Olive White-eye (green), and Mauritius Grey White-eye (orange). Grey colours represent elevation classes; for Reunion: light grey = elevation below 1,000 m, medium grey = elevation between 1,000 and 2,000 m, dark grey = elevation above 2,000 m; for Mauritius: light grey = elevation below 400 m, medium grey = elevation between 400 and 600 m, dark grey = elevation above 600 m. Dotted lines separate the distribution ranges of the four Reunion Grey White-eye geographic forms. Phylogenetic relationships between the four endemic Mascarene white-eyes are adapted from Warren et al. (2006), with the asterisk denoting the divergence time between the two sister lineages, estimated at 1.2 Mya. Illustrations from Birds of the World reproduced with permission from Lynx edicions. Photo credits: Stanislas Harvančík (Reunion Grey White-eye), Simon Colenutt (Reunion Olive White-eye), and Hilary Jones (Mauritius Grey White-eye).

## Materials and Methods

### a) Population sampling and DNA sequence data

Blood samples from a total of 42 individuals, captured in mistnets and then released, were collected between 2007 and 2017 throughout the entire distribution of the Reunion Grey White-eye (22 individuals from 18 localities), the Reunion Olive White-eye (11 individuals from 11 localities), and the Mauritius Grey White-eye (nine individuals from nine localities) (Figure 2; Table S1). Approximately 1 µg of high-quality DNA was extracted using a QIAGEN DNeasy Blood & Tissue kit following the manufacturer’s instructions, with an extra pre-digestion grinding step. Genomic DNA extractions were sent to Novogene Bioinformatics Technology (Beijing, China; 25 individuals) and GeT-PlaGe core facility (INRA, Toulouse; nine individuals) for shotgun whole-genome resequencing with an Illumina HiSeq2500 sequencer. Our dataset includes data from eight individuals sequenced using similar procedures in previous studies (Bourgeois et al. 2017; Leroy, Anselmetti, et al. 2021).

### b) Reference-based whole-genome processing

To genotype individuals, sequencing reads of all three species were first mapped against the high-quality genome assembly (scaffold N50 exceeding one megabase) of a Reunion Grey White-eye (Leroy, Anselmetti, et al. 2021) using the bwa-mem algorithm (Li and Durbin 2009) with default parameters. Single nucleotide polymorphisms (SNPs) were then called for each species independently using GATK v. 3.7 (McKenna et al. 2010) using base quality score recalibration and indel realignment. Both variant and invariant sites were called (option “--includeNonVariantSites”). Variant filtration was performed using a custom script to speed up computations, but following the same procedures than under GATK, using the following thresholds: QD>2.0, FS<60.0, MQ>40.0, MQRANKSUM>-2.0, READPOSRANKSUM>-2.0 and RAW_MQ > 45,000. The three species were also genotyped together for multi-population demographic inferences, using the same variant filtration procedure. After filtering, the mean individual coverage was 15x (see Table S2 for details about the samples used). To check for the general consistency of the obtained genomic dataset, we performed a Principal Component Analysis with PLINK v. 1.90b5.3 (Purcell et al. 2007) and plotted the results in R version 3.5 (R core team 2020) with adegenet v. 2.1.1 (Jombart 2008) (Figure S1).

### c) Mutation rate and generation time estimates

Mutation rate and generation time are two key parameters in population genetic inference. In this study, we used the same set of parameter values for the three focal species, considering that they diverged from a common ancestor 1.2 million years ago at most (Martins et al. 2020; Warren et al. 2006), and genomic features tend to be conserved in birds (Kawakami et al. 2014; Singhal et al. 2015). We used a mutation rate of 4.6e^-9^ mutations per site per generation (95% confidence interval: 3.4e^-9^ - 5.9e^-9^) based on a direct estimate for the Collared Flycatcher (*Ficedula albicollis* (Temminck, 1815)) (Smeds, Qvarnström, and Ellegren 2016). A few estimates of generation times are available for *Zosterops* species, suggesting short generation times, possibly less than a year (Cornetti et al. 2015; Moyle et al. 2009), but generation time is between two and three years in the Heron Island population of Silvereye (*Zosterops lateralis chlorocephalus* Gould, 1841) (Clegg et al. 2002; Estoup and Clegg 2003). Thus, we used a generation time of one year but also run our analyses using values ranging between 0.5 and 2 years to account for uncertainty.

### d) Demographic inferences

We used two independent approaches to reconstruct the demographic histories of the three Mascarene white-eyes.

#### Markovian coalescent analyses

In order to estimate the historical population size trajectory in each species, we first performed inferences of *N_e_* using the Pairwise Sequentially Markovian Coalescent (PSMC, Li and Durbin, 2011) since this method can be readily applied to unphased whole-genome sequence data. Briefly, the PSMC model infers the local time to the most recent common ancestor (TMRCA) on the basis of the local density of heterozygotes and estimates the inverse instantaneous rate (IICR) of a sample which is equivalent to population size in panmictic models (Mazet et al. 2016). We considered two different thresholds for the minimum sequence length, either 1,000 bp or 100,000 bp. For each individual, a custom script was used to convert genotypes into PSMC input format, *i.e.* converting each 100-base-pair window into K if the window contains at least one heterozygote site and T otherwise, using the following filters: QUAL > 100, 10<depth<150, N_ratio (ratio of missing data in the window) < 0.9. Sites outside these thresholds were considered to be missing (N). We set the upper limit of the TMRCA to 5 and the initial θ/ρ value to 1 and we used the atomic interval scheme “4+30*2+4+6+10”, as in Nadachowska-Brzyska et al. (2016).

#### Approximate Bayesian Computation inferences

We performed ABC inferences from whole-genome sequence data using DILS (Demographic Inferences with Linked Selection; Fraïsse et al., 2021). This pipeline gathers ABC demographic inferences of population size changes and timing of gene flow during divergence using either single-population models (Figure S2A) or two-population models (Figure S2B). As DILS needs sequences as input, we performed a whole-genome sequence reconstruction at the individual level (two sequences per chromosome considering diploidy) for all individuals as published in Leroy, Rousselle, et al. (2021), and then generated DNA segments of 10 kb for all the scaffolds with a scaffold size exceeding 10 kb and kept scaffolds between 1 and 10 kb as one DNA segment. In addition, reconstructed sequences with an N content exceeding 50% (missing information) were discarded. Eight summary statistics were used for single-population models: number of biallelic sites, pairwise nucleotide diversity (π) (Tajima 1983), Watterson’s *θ* (Watterson 1975) and Tajima’s *D* (Tajima 1989), considering the average and standard deviation of each measure. Thirty-seven summary statistics were used for two-population models, including the same estimates of genetic diversity for each population, and measures of differentiation and divergence between populations (F_ST_; B. S. Weir & Cockerham, 1984; Wright, 1943); net divergence; Nei and Li, 1979) (see Table S3 for a list of all summary statistics). For all inferences, model comparison was performed using a random forest algorithm. Two million simulations were run under the best model to perform parameter estimations, and four rounds of goodness-of-fit were run to refine parameter estimation (Fraïsse et al. 2021).

We first used single-population models for each of the three species to compare three scenarios involving either a constant population size (i) or a change of population size over time, with either an expansion (ii) or a contraction (iii) (Figure S2A). Models including homogeneous or heterogeneous *N_e_* across the genome were compared. Three parameters were estimated for single-population models, namely population size prior to the population size change event (N_ANC_), population size after this event (N_POP_), and time since this event in number of generations (T_dem_). The priors used for each parameter are detailed in Table S4.

Since single-population models assume closed systems, we also used models of two-population divergence with different timing of gene flow, with two models assuming current isolation, *i.e.* absence of ongoing gene flow, (SI: Strict Isolation; AM: Ancestral Migration), and two models assuming ongoing gene flow (IM: Isolation with Migration; SC: Secondary Contact) (Figure S2B). These four models assume the instantaneous split of an ancestral population into two daughter populations that diverge either without any migration (SI), with continuous introgression since the time of split (IM), with a period of migration in the early times of lineage divergence only (AM), or with a period of migration occurring upon secondary contact after an initial period of strict isolation (SC). The models also included a possible change in population size in the daughter populations after the ancestral split. Eight parameters were estimated for all two-population models, namely the population size of the ancestral population (N_ANC_), the time of the split between the two populations (T_split_), the population sizes of the daughter populations after the split (N_ANC1_ and N_ANC2_), the time of the potential demographic change (T_dem1_ and T_dem2_) and the population sizes of the daughter populations after the demographic change (N_POP1_ and N_POP2_). The time of the secondary contact (T_SC_) was estimated in the SC model, and the time of ancestral migration cessation (T_AM_) was estimated in the AM model. Daughter population migration rates were estimated for the three models involving migration (IM, AM, SC). The priors used for each parameter are detailed in Table S4. We run these models for the three possible pairs of species (Reunion Grey White-eye - Mauritius Grey White-eye; Reunion Grey White-eye - Reunion Olive White-eye and Mauritius Grey White-eye - Reunion Olive White-eye).

## Results

Population size history reconstructions though PSMC analyses support contrasting population size trajectories between species living on different islands (Figure 3). While cycles of expansions and contractions are observed in the Reunion Grey and Olive White-eyes, Mauritius Grey White-eyes present a clear signal of long-term expansion. Specifically, in the Reunion Grey White-eye, a population decline was detected that could coincide with the Last Glacial Maximum (LGM; 21,000 ya), while, over the same period of time, only a slight reduction or population size stasis was detected in the Reunion Olive White-eye. In contrast, PSMC analyses indicate that the Mauritius Grey White-eye underwent an expansion in the corresponding time window, suggesting that the LGM might have had a lower impact on this species than on the Reunion species. Similar demographic histories were reconstructed for the different geographic forms of the Reunion Grey White-eye (Figure S3), with a possibly stronger reduction in population size around the time of the LGM in the highland form. Even after considering substantial variation in the mutation rate and generation time (Figure S4), our main conclusions still hold. Similarly, restricting the analysis to the longest sequences with a minimal length of 100,000 bp instead of 1,000 bp led to similar results (Figure S5), suggesting that our results are robust to a certain extent to methodological limits and parameter uncertainties.

**Figure 3:**
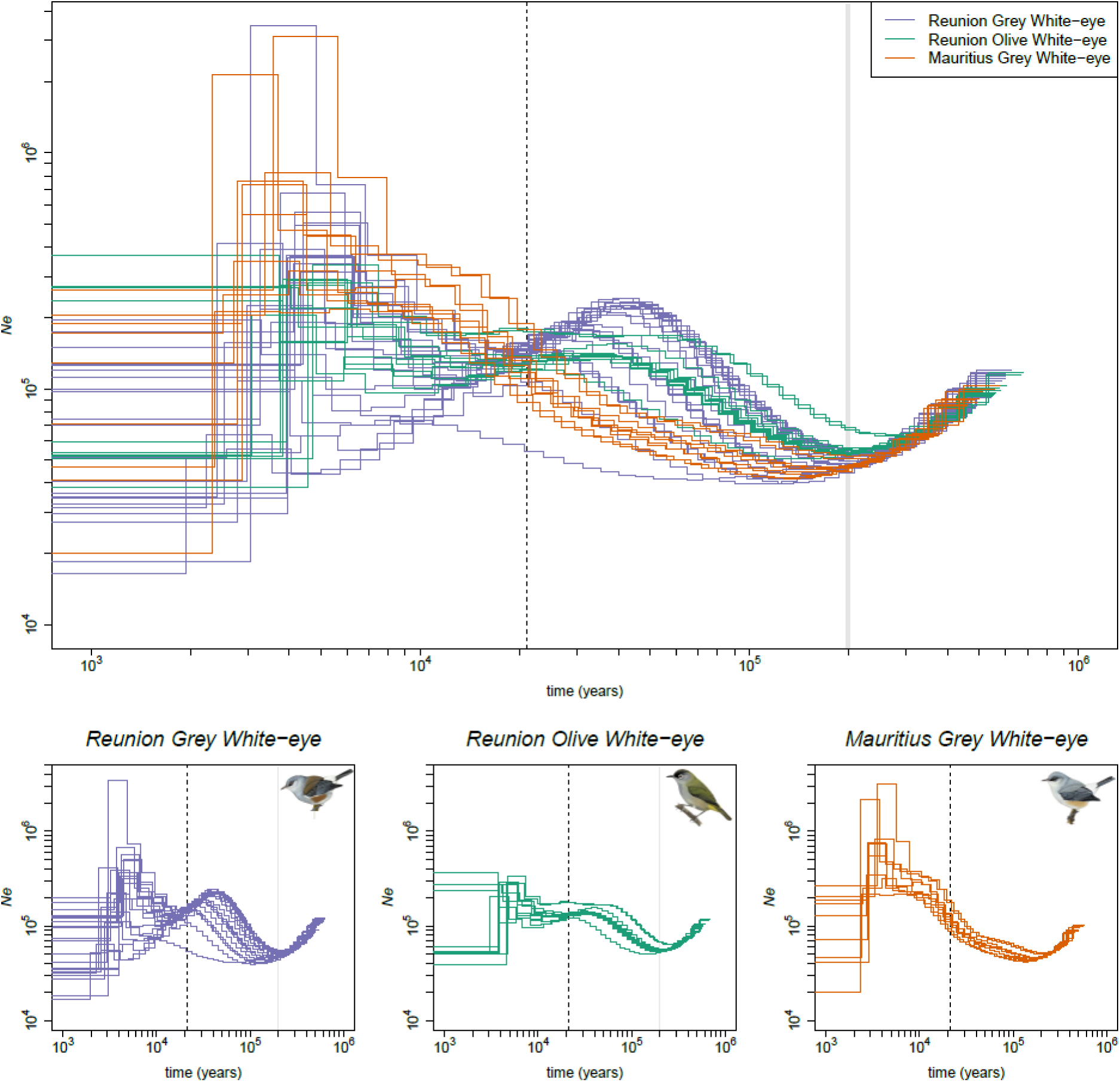
PSMC inferences of population size histories over time. Colours correspond to the three white-eye species, and each line represents a different individual. Vertical dotted lines and grey vertical bars indicate (from left to right): the Last Glacial Maximum 21,000 ya and a catastrophic period of large explosive eruptions in Reunion around 200,000 ya. In Mauritius, volcanic activity has been quite intense throughout the last 500,000 years but was never explosive during that period. Please note that the plots are in log-scales. Illustrations from Birds of the World reproduced with permission from Lynx edicions.

In general agreement with the PSMC analyses, ABC demographic reconstruction highlighted a strong signal of expansion for all species, in single-population models as well as in two-population models. For single-population inferences, models including an expansion phase had a very high posterior probability (≥ 0.95) compared to models including a bottleneck or constant population size (Table S5). More specifically, ancestral *N_e_* values were estimated to be around 35,000, 65,000 and 130,000 individuals for the Reunion Grey White-eye, Mauritius Grey White-eye and Reunion Olive White-eye, respectively, and current *N_e_* values were estimated to be around 100,000, 490,000 and 700,000, respectively, representing 2.8-, 7.5- and 5.4-fold increases (Table S6). Based on these models, the onsets of expansion were very different between Reunion and Mauritius Grey White-eyes (around 20,000 generations ago and 45,000 generations ago, respectively) and the Reunion Olive White-eye (around 250,000 generations ago).

Two-population models of divergence inferred an Ancestral Migration scenario as the most likely (posterior probability higher than 0.81 for current isolation models and higher than 0.67 for Ancestral Migration versus Strict Isolation; Figure S6), with a brief period of ancestral gene flow in all species comparisons (median values for the T_AM_/T_SPLIT_ ratio: 0.72, 0.94 and 0.73 for Reunion Grey White-eye - Mauritius Grey White-eye; Reunion Grey White-eye - Reunion Olive White-eye and Mauritius Grey White-eye - Reunion Olive White-eye, respectively). Evidence for current isolation between species was expected, especially for species living in different islands and estimates of divergence times that were obtained in our models are consistent with previous estimates (Cai et al. 2019; Warren et al. 2006), suggesting that the temporal framework of our demographic inferences using ABC may be quite robust. Parameter estimations obtained using these models suggested 10-fold expansions from ancestral *N_e_* of between 35,000 and 75,000 individuals to current *N_e_* between 400,000 and 800,000 individuals across the three species (Table 1). These expansions seemed to be rather ancient for the Reunion Grey and Olive White-eyes, with an expansion estimated around 70-90% of the time of divergence, while the expansion inferred for the Mauritius Grey White-eye seemed more recent (30-50% of the time of divergence), highlighting again the importance of island-specific events on population size trajectories.

**Table 1:** Parameter estimation for two-population demographic scenarios in DILS. Numbers outside brackets indicate median values while numbers inside brackets indicate confidence intervals (2.5%-97.5%). Nine parameters were estimated, namely the population size of the ancestral population (N_ANC_), the time of the split between the two populations (T_split_), the population sizes of the daughter populations after the split (N_ANC1_ and N_ANC2_), the time of the potential demographic change (T_dem1_ and T_dem2_), the population sizes of the daughter populations after the demographic change (N_POP1_ and N_POP2_), and the time of ancestral migration cessation (T_AM_). For each pair, populations 1 and 2 are the first and second population of the pair, respectively. See Figure S2B for a representation of this scenario of divergence.

|  | ABC model parameters |  |  |  |  |  |  |  |  |
| --- | --- | --- | --- | --- | --- | --- | --- | --- | --- |
| | $N_{ANC}$ | $T_{split}$ | $N_{ANC1}$ | $N_{ANC2}$ | $T_{dem1}$ | $N_{POP1}$ | $T_{dem2}$ | $N_{POP2}$ | $T_{AM}$ |
| *Reunion Grey White-eye - Mauritius Grey White-eye* | 115,681<br>(103,039-130,232) | 128,291<br>(95,638-168,209) | 46,070<br>(24,675-76,275) | 48,886<br>(30,636-77,610) | 98,879<br>(80,019-121,191) | 631,186<br>(487,328-742,422) | 69,215<br>(55,562-86,687) | 537,411<br>(396,932-661,929) | 92,045<br>(75,532-110,466) |
| *Reunion Grey White-eye (GHB form) - Mauritius Grey White-eye* | 84,075<br>(73,819-97,160) | 115,323<br>(86,075-152,274) | 66,184<br>(36,219-109,992) | 47,120<br>(27,039-74,713) | 91,004<br>(69,428-116,346) | 443,404<br>(338,203-581,931) | 60,632<br>(46,932-80,056) | 405,894<br>(284,879-536,721) | 93,817<br>(74,518-112,903) |
| *Reunion Grey White-eye - Reunion Olive White-eye* | 234,347<br>(200,173-275,027) | 318,314<br>(255,817-398,370) | 67,133<br>(22,956-120,907) | 55,254<br>(24,862-96,435) | 234,291<br>(188,145-290,008) | 797,268<br>(604,018-915,561) | 293,743<br>(234,835-364,791) | 816,811<br>(651,688-945,258) | 299,302<br>(241,213-350,777) |
| *Mauritius Grey White-eye - Reunion Olive White-eye* | 237,298<br>(192,706-277,644) | 431,242<br>(299,381-559,243) | 76,860<br>(41,129-120,390) | 34,639<br>(16,827-58,066) | 118,964<br>(91,229-141,605) | 745,166<br>(556,712-916,230) | 291,374<br>(218,052-356,087) | 407,800<br>(322,593-453,945) | 316,104<br>(258,245-385,771) |

Results from ABC demographic inferences for one of the four geographic forms of the Reunion Grey White-eye, the Grey-Headed Brown form (GHB), are consistent with those considering all Reunion Grey White-eye geographic forms together (Tables S5 and S6, Table 1), suggesting that the different geographic forms may have experienced similar population trajectories.

## Discussion

Reconstructing long-term population dynamics using whole-genome data has been key in revealing how past regional climatic events influenced changes in population size through time in many different taxa (*e.g.* Chattopadhyay et al., 2019; Germain et al., 2023; Nadachowska-Brzyska et al., 2015). Yet, populations and species face ecological disruptions over time that can vary from being temporary and localized to regional or even global with no necessary synchrony between the two types of events. Therefore, estimates of the changes in effective population sizes as visualized in PSMC profiles may reflect demographic responses to a number of distinct events acting at different temporal and spatial scales and individual effects that may be difficult to distinguish from one another. In addition, the combined effects of these different sources of variation on population demographic histories may also be accompanied by variation in population structure, to which current approaches have been shown to be very sensistive (Chikhi et al. 2018; Mazet et al. 2016; Wakeley 1999). In an attempt to alleviate this difficulty and distinguish the effects of local and regional disruption events on population demographic trajectories, our study used volcanic oceanic islands as a spatial and temporal framework, whole-genome data obtained from closely related species, and ABC demographic inference in addition to PSMC to uncover population size changes that could be attributable to the different types of events. We obtained two key findings. First, we found that species from the same island (Reunion) showed similarities in their population size trajectories that differed from the one reconstructed for the species from the other island (Mauritius); and second, that this pattern also applied to within-island geographic forms of the Reunion Grey White-eye that exhibited similar demographic histories, distinct from the demographic history of the sister taxon, the Mauritius Grey White-eye.

These findings suggest that local events have played a significant role in shaping population trajectories in these island bird species. They also highlight the potential of our conceptual framework and its working hypotheses in disentangling the effects of local and regional drivers on past demography of species. This is crucial to obtain a better understanding of long-term population processes that are of paramount conservation importance in the current context of rapid environmental change driven by both local and global factors of ecological disruption (Leung et al. 2020; Nogués-Bravo et al. 2018; Urban et al. 2016).

What could be the factors explaining shared response to local (island-specific) ecological disruptions in our study system? In the two Reunion species of white-eyes, the period of lowest *N_e_* corresponds approximately to the LGM, while in the Mauritius Grey White-eye, this episode seems to have had no effect on the demographic history. Thus, global drivers like LGM seem to have some effects but the response to these effects may vary depending on local conditions and the ecology of the species, although this last factor is probably unimportant here given how closely related and ecologically similar white-eye species are in both islands (Cheke 1987). Similar evidence of a bottleneck associated to the LGM timing has been observed in the Reunion coffee tree (*Coffea mauritiana* Lam., 1783) (Garot et al. 2019) and was interpreted as being the result of a range contraction due to major drops in both precipitation (Yan, Wang, and Liu 2016) and temperature (Barrows and Juggins 2005), especially on the leeward side of the island. Since the Reunion coffee tree is a widespread forest species, this may imply that forest extent may have been greatly reduced during the LGM, which might have impacted bird populations, especially at lower elevations, according to a well-known scenario of distribution shifts in response to Quaternary climatic fluctuations (Hewitt 2000). Reduction of forest area at the LGM and restriction in refugia have been reconstructed in many instances (*e.g.* Carnaval and Moritz, 2008) with possible demographic consequences on the fauna the forest remnants sheltered (*e.g.* Maldonado-Coelho, 2012). Elevational downward shifts of ecosystems towards the coast can be expected during glacial events (Fernández-Palacios et al. 2016), especially in high-elevation islands like Reunion (see Salces-Castellano et al., 2021). The highland habitat might have become less suitable and the species’ range may have contracted towards the coast. In line with this, our PSMC results indicate that the highland form of the Reunion Grey White-eye may have experienced a stronger bottleneck associated with the LGM than the other forms restricted to the lowlands. By contrast, our Mauritius Grey White-eye PSMC analyses suggest that the population kept expanding during the LGM. As Mauritius is a low elevation island, it is possible that during the LGM Mauritius offered more suitable lowland habitats compared to Reunion. Furthermore, the globally low sea level associated with the LGM translated into an increase in land area that was more substantial for Mauritius (50%) than for Reunion (10%) (Norder et al. 2018). The 4,200 BP megadrought that has been documented in East Africa (Marchant and Hooghiemstra 2004; Thompson 2002), North America (Booth et al. 2005), India (Staubwasser et al. 2003), Madagascar (Gasse and Van Campo 2001) and Mauritius (de Boer et al. 2015; Rijsdijk et al. 2011) and has been associated with massive mortality and population bottlenecks of Western Indian ocean and African species (Heller et al. 2008; Rijsdijk et al. 2011; Salmona et al. 2012; Salmona, Heller, Lascoux, et al. 2017) must have impacted populations of our study species, but it left no detectable signature in the population trajectories that we recovered. However, PSMC analysis lacks resolution at such recent dates, as shown in Figure 3, and spurious signals of bottlenecks are often observed in the lower time limit of PSMC efficiency (1,000 - 10,000 ya) limiting the use of that approach for recent periods (Li and Durbin 2011). While some new methods have become available to reconstruct demographic histories close to present (Terhorst et al. 2017), understanding the impact on population size of very recent events still remains a challenge and requires larger datasets with multiple resequenced individuals per species (Patton et al. 2019; Wilder et al. 2023). Lastly, we did not detect bottlenecks associated with major dated volcanic eruptions that occurred in either Reunion or Mauritius, as it has been observed in other island bird systems (Bemmels et al. 2022). This could indicate that the volcanic eruptions in the Mascarenes did not affect the totality of the islands in spite of the magnitude of some events, in particular a series that occurred around 200 kya on Reunion and are thought to have wreaked havoc on a large area of the island (Castellanos Melendez et al. 2023; Fretzdorff et al. 2000), and/or that the white-eyes were able to find habitat refugia in which they maintained relatively high population sizes in spite of the possible devastation of most of the island. This view is supported by our results from both PSMC and ABC demographic inferences that strongly support ancient and long-term expansions in the three study species, likely associated with evolutionary processes which took place on the islands themselves since the species diverged from an ancestral stock and reflecting the radiation of the Mascarene white-eyes (Warren et al. 2006).

While our study recovered potentially important information about how the demographic history of island species may have been influenced by ecological disruption acting at different spatial and temporal scales, it also highlights a series of difficulties and limitations inherent to this kind of investigations. First, population structure was not explicitly taken into account in our study; yet, it can severely bias demographic inferences. For instance, it has been shown that a PSMC signal consistent with population size changes can be obtained in a population without any change of population in its history but with just changes in connectivity between subpopulations (Grusea et al. 2019; Mazet et al. 2016; Rodríguez et al. 2018). Our PSMC reconstructions were similar for all four geographic forms of the Reunion Grey White-eye, and all ABC inferences for one particular geographic form (the GHB form) produced similar results to those obtained using all geographic forms together, making us confident that our conclusions are fairly robust to deviations from panmixia due to population structure, at least for that species. Second, we assumed similar generation times for the three species, while these have different social structure and population densities, that may influence this parameter. For example, cooperative breeding has been reported in Reunion and Mauritius Grey White-eyes (Gill 1973; Hansen, Olesen, and Jones 2002) that may delay first reproduction and overall increase generation time (Kreider et al. 2022). However, we note that the results from PSMC analyses did not change substantially after varying both mutation rate and generation time, suggesting that our results are also robust, to some extent, to variations in these parameters. Third, we only considered models with up to two populations since the framework that we used (DILS; Fraïsse et al., 2021), while fast and malleable, only works with two populations at most. We tried using the Fastsimcoal modelling approach (Excoffier et al. 2013) as an alternative but when compared to DILS, it yielded unrealistic results with very poor fits to the data (results not shown). In addition, Fastsimcoal does not allow for *N_e_* and N_m_ heterogeneity along the genome, which can lead to inaccurate demographic inferences while DILS parametrizes this heterogeneity. Our ABC analyses indeed unconditionally support *N_e_* heterogeneity along the genome (posterior probabilities of 0.99 for all the species, Table S4) indicating that accounting for this factor may be critical in demographic inference based on whole-genome sequence data. While strong signals of expansion were detected in our ABC demographic inferences, we cannot really rule out the possibility that bottlenecks occurred over the course of the species’ population histories. More complex ABC models including several waves of demographic changes would be necessary to properly address this, but current ABC approaches may not be powerful enough to detect ancient and subtle population size changes. However, PSMC analyses, which account for the possibility of multiple population size changes across time, do not suggest many demographic changes, but are rather consistent with a general strong expansion.

To conclude, the shared population histories in the Reunion Grey and Olive White-eyes and their contrast with the Mauritius Grey White-eye population history suggest that, in this system and at a relatively small spatial scale, local events may have been important drivers of population trajectories. Using more species from Reunion and Mauritius will be necessary to further validate whether the differences in demographic trajectories that we recovered are general. As white-eyes could be more resilient to environmental changes than other species groups, the use of other species, with different ecologies or being less behaviourally plastic, could yield different population trajectories in response to both local and regional events. More importantly, investigating the demographic history of multiple species from a range of taxa in several archipelagos by comparing species living on the same or different islands and controlling for their evolutionary relatedness, as recently advocated by Cerca et al. (2023) for another purpose, would greatly help understanding how species respond to local and regional events of ecological disruption and would provide an useful tool for assessing the impacts and risks of plausible future human-driven changes in climate.

## Acknowledgments

We thank Thomas Duval, Jennifer Devillechabrolle, Guillaume Gélinaud, Marie Manceau, Juli Broggi, Josselin Cornuault, Yann Bourgeois, Joris Bertrand, Boris Delahaie, Dominique Strasberg, Philipp Heeb, René-Claude Billot, Ben Warren and Jean-Michel Probst for their assistance in the field in Reunion and Mauritius; Reunion National Park and Mauritius National Parks and Conservation Service for granting us permission to conduct fieldwork in protected areas of Reunion and Mauritius, respectively; Benoit Lequette (Reunion National Park) and Vikash Tatayah (Mauritius Wildlife Foundation) for facilitating permit collection. We thank Hélène Holota, Uxue Suescun, and Amaya Iribar-Pelozuelo for help with the laboratory work; Lounès Chikhi for stimulating discussions on population structure and demographic inferences; Camille Roux for his valuable advice on the use of DILS for demographic inferences; the IFB cluster and Genotoul bioinformatics platform for the computational resources they provided and in particular Marie-Stéphane Trotard for bioinformatics support. We are also grateful to Stanislav Harvančík, Simon Colenutt, and Hilary Jones for allowing us to reproduce their photos of white-eyes.

## Data Accessibility and Benefit-Sharing

Raw sequence reads of whole genomes are deposited in the SRA (BioProject PRJNA661201 and PRJEB18566). See Table S2 for accession codes of all the samples used in this study.

## Author Contributions

B.M. and C.T. initiated and coordinated the project. M.G., B.N., B.M. and C.T. conceived the study and designed the experiments. B.M. and C.T. conducted the fieldwork, with the assistance of M.G., T.L. and B.N. M.G. performed the analyses. M.G., T.L., J.S., B.N., B.M. and C.T. interpreted the analyses and wrote the manuscript.

