## Supplemental Information for "Demographic responses of oceanic island birds to local and regional ecological disruptions revealed by whole-genome sequencing"

Maëva Gabrielli, Thibault Leroy, Jordi Salmona, Benoit Nabholz, Borja Milá, Christophe Thébaud

**Table of Contents:**

| **Details about sampling localities** | Page 2 |
| --- | --- |
| **Details about samples** | Pages 3,4 |
| **Summary statistics for DILS** | Page 5 |
| **Priors used in DILS** | Page 6 |
| **Model choice for single population inferences in DILS** | Page 7 |
| **Parameter estimation for single population inferences in DILS** | Page 8 |
| **PCA of SNPs from all individuals** | Page 9 |
| **Schematic representations of all models used in DILS** | Page 10 |
| **PSMC of the different geographic forms of the Reunion Grey White-eye** | Page 11 |
| **PSMC with different generation times and mutation rates** | Page 12 |
| **PSMC with a minimal length of 1,000 pb or 100,000 pb** | Page 13 |
| **Schematic view of the DILS pipeline for two population models** | Page 14 |

**Table S1**. Details about the 33 localities used for genomic analyses, with corresponding locality number used in Figure 2, including the region, latitude, longitude, and elevation in meters of each sampling locality.

| Sampling site | Region | Latitude | Longitude | Elevation (m) |
| --- | --- | --- | --- | --- |
| 1/ Grande Chaloupe | Reunion | -20.9005 | 55.3758 | 35 |
| 2/ Moka | Reunion | -20.9279 | 55.5150 | 219 |
| 3/ Rivière du Mât les Bas | Reunion | -20.9794 | 55.6823 | 42 |
| 4/ Sentier Ste-Marguerite | Reunion | -21.1088 | 55.6883 | 550 |
| 5/ Trois Citernes | Reunion | -21.1589 | 55.7618 | 846 |
| 6/ Coulée 2007 | Reunion | -21.2782 | 55.7916 | 194 |
| 7/ Basse Vallée | Reunion | -21.3409 | 55.7090 | 687 |
| 8/ Piton de l'Entonnoir | Reunion | -21.3566 | 55.6145 | 430 |
| 9/ Rivière d'Abord | Reunion | -21.3254 | 55.4949 | 160 |
| 10/ L'Etang-salé les Bains | Reunion | -21.2585 | 55.3379 | 68 |
| 11/ St-Leu | Reunion | -21.1374 | 55.2953 | 509 |
| 12/ Ermitage | Reunion | -21.0733 | 55.2327 | 60 |
| 13/ Petit Bernica | Reunion | -21.0254 | 55.2803 | 306 |
| 14/ Maïdo | Reunion | -21.0728 | 55.3774 | 2,062 |
| 15/ Tévelave | Reunion | -21.1689 | 55.3871 | 1,962 |
| 16/ Bébour | Reunion | -21.0965 | 55.5495 | 1,540 |
| 17/ Bois Ozoux | Reunion | -21.1979 | 55.6465 | 2,283 |
| 18/ Pas de Bellecombe | Reunion | -21.2172 | 55.6877 | 2,246 |
| 19/ Anse des Cascades | Reunion | -21.1837 | 55.8301 | 55 |
| 20/ Canot | Reunion | -21.2240 | 55.4131 | 690 |
| 21/ Roche Verre Bouteille | Reunion | -20.9877 | 55.3951 | 1,251 |
| 22/ Grand Matarum | Reunion | -21.1244 | 55.4787 | 1,460 |
| 23/ Nez de Bœuf | Reunion | -21.2062 | 55.6189 | 2,070 |
| 24/ Ravine Petit St-Pierre | Reunion | -21.1850 | 55.6722 | 2,076 |
| 25/ Cap Malheureux | Mauritius | -19.9893 | 57.6284 | 21 |
| 26/ Roches Noires forest | Mauritius | -20.1153 | 57.7357 | 15 |
| 27/ Le Bouchon | Mauritius | -20.4709 | 57.6786 | 15 |
| 28/ Bel Ombre Forest | Mauritius | -20.4735 | 57.4207 | 278 |
| 29/ Le Morne Brabant | Mauritius | -20.4598 | 57.3323 | 9 |
| 30/ Black River Gorges | Mauritius | -20.3836 | 57.4199 | 117 |
| 31/ Macchabé–Brise Fer forest | Mauritius | -20.3787 | 57.4407 | 600 |
| 32/ Yemen | Mauritius | -20.3432 | 57.4138 | 164 |
| 33/ Le Pouce Mt | Mauritius | -20.1990 | 57.5238 | 605 |

**Table S2**. Details about the samples used in our study, including the locality, mean individual depth of coverage after SNP calling and sample accession code from the ENA platform.

| **Species** | **Individual code** | **Locality** | **Mean depth** | **ENA sample accession** |
| --- | --- | --- | --- | --- |
| Reunion Grey White-eye | 1243 | Tévelave | 19.68 | SAMN16057078 |
| Reunion Grey White-eye | 1544 | Moka | 12.39 | SAMN16057079 |
| Reunion Grey White-eye | 1681 | Rivière d'Abord | 9.48 | SAMN16057080 |
| Reunion Grey White-eye | 2062 | Rivière du Mât les Bas | 18.60 | SAMN16057081 |
| Reunion Grey White-eye | 2078 | Sentier Ste-Marguerite | 16.66 | SAMN16057082 |
| Reunion Grey White-eye | 487 | St-Leu | 17.83 | SAMN16057076 |
| Reunion Grey White-eye | 604 | L'Etang-salé les Bains | 13.63 | SAMN16057077 |
| Reunion Grey White-eye | 314 | Maïdo | 9.19 | SAMEA24914668 |
| Reunion Grey White-eye | 317 | Maïdo | 9.91 | SAMEA24915418 |
| Reunion Grey White-eye | 1430 | Pas de Bellecombe | 10.79 | SAMEA24920668 |
| Reunion Grey White-eye | 1434 | Pas de Bellecombe | 9.17 | SAMEA24922168 |
| Reunion Grey White-eye | 1337 | Bois Ozoux | 8.17 | SAMEA24926668 |
| Reunion Grey White-eye | 1588 | Bois Ozoux | 6.40 | SAMEA24928168 |
| Reunion Grey White-eye | JB41 | Grande Chaloupe | 13.91 | SAMN16057086 |
| Reunion Grey White-eye | JB58 | Petit Bernica | 25.88 | SAMN16057087 |
| Reunion Grey White-eye | 11_0809 | Trois Citernes | 20.03 | SAMN16048776 |
| Reunion Grey White-eye | 11_1020 | Coulée 2007 | 10.91 | SAMN16049245 |
| Reunion Grey White-eye | 1602 | Piton de l'Entonnoir | 10.84 | SAMN16049248 |
| Reunion Grey White-eye | 17_670 | Bébour | 17.28 | SAMN16049249 |
| Reunion Grey White-eye | 264 | Basse Vallée | 15.83 | SAMN16049246 |
| Reunion Grey White-eye | 996 | Ermitage | 11.10 | SAMN16049247 |
| Reunion Grey White-eye | 15179 | Pas de Bellecombe | 38.71 | SAMN11339519 |
| Reunion Olive White-eye | 1113 | Sentier Ste-Marguerite | 19.29 | SAMN16057384 |
| Reunion Olive White-eye | 11-1034 | Anse des Cascades | 10.96 | SAMN16057375 |
| Reunion Olive White-eye | 1418 | Canot | 10.67 | SAMN16057377 |
| Reunion Olive White-eye | 1550 | Moka | 11.37 | SAMN16057382 |
| Reunion Olive White-eye | 1649 | Bébour | 10.54 | SAMN16057376 |
| Reunion Olive White-eye | 17-691 | Grand Matarum | 11.14 | SAMN16057380 |
| Reunion Olive White-eye | 2217 | Nez de Bœuf | 12.33 | SAMN16057383 |
| Reunion Olive White-eye | 236 | Basse Vallée | 12.05 | SAMN16057379 |
| Reunion Olive White-eye | 322 | Maïdo | 10.56 | SAMN16057381 |
| Reunion Olive White-eye | 354 | Roche Verre Bouteille | 8.87 | SAMN16057378 |
| Reunion Olive White-eye | 15-026 | Ravine Petit St-Pierre | 52.55 | SAMN16057409 |
| Mauritius Grey White-eye | 1281 | Le Bouchon | 10.59 | SAMN16057367 |
| Mauritius Grey White-eye | 1295 | Le Morne Brabant | 12.76 | SAMN16057374 |
| Mauritius Grey White-eye | 1310 | Roches Noires forest | 11.53 | SAMN16057369 |
| Mauritius Grey White-eye | 1312 | Cap Malheureux | 11.02 | SAMN16057368 |
| Mauritius Grey White-eye | 1319 | Le Pouce Mt | 20.46 | SAMN16057373 |
| Mauritius Grey White-eye | 1326 | Macchabé–Brise Fer forest | 12.24 | SAMN16057371 |
| Mauritius Grey White-eye | 391 | Yemen | 12.63 | SAMN16057370 |
| Mauritius Grey White-eye | 398 | Black River Gorges | 16.68 | SAMN16057372 |
| Mauritius Grey White-eye | 461 | Bel Ombre Forest | 10.68 | SAMN16057366 |

**Table S3**. Summary statistics used for two-population demographic inferences in DILS. * indicate that the average and standard variation of the statistics were used as two different summary statistics

| **statistics** | **description** |
| --- | --- |
| bialsites * | total number of biallelic sites |
| *Sf* * | number of fixed differences between both species |
| *Sx_A_* * | number of biallelic positions exclusively polymorphic in spA |
| *Sx_B_* * | number of biallelic positions exclusively polymorphic in spB |
| *Ss* * | number of biallelic positions exclusively polymorphic shared by both species |
| *π_A_* * | nucleotide *π* in species A |
| *π_B_* * | nucleotide *π* in species B |
| pearson_r_pi | Pearson's R² correlation coefficient in nucleotide *π* |
| *θ_A_* * | Watterson's *θ* in species A |
| *θ_B_* * | Watterson's *θ* in species B |
| pearson_r_theta | Pearson's R² correlation coefficient in *θ_W_* |
| DtajA * | Tajima's *D* in species A |
| DtajB * | Tajima's *D* in species B |
| divAB * | total interspecific divergence |
| netdivAB * | net interspecific divergence |
| minDivAB * | minimal interspecific divergence |
| maxDivAB * | maximal interspecific divergence |
| FST * | Fixation index (*F_ST_*) between species A and species B |
| pearson_r_divAB_netDivAB | Pearson's R² correlation coefficient between divAB and netDivAB |
| pearson_r_divAB_FST | Pearson's R² correlation coefficient between divAB and *F_ST_* |
| pearson_r_netdivAB_FST | Pearson's R² correlation coefficient between netdivAB and *F_ST_* |

**Table S4**. Priors for demographic inferences in DILS: mutation rate (mu), ratio of population recombination over population mutation (rho_over_theta), minimum and maximum population size in coalescent units (N_min and N_max), minimum and maximum time of demographic change in coalescent units (Tchanges_min and Tchanges_max), minimum and maximum time of split in coalescent units (Tsplit_min and Tsplit_max) and minimum and maximum migration rate in number of migrant per generation (M_min and M_max). * indicate the priors only needed for single-population analyses while ** indicate the priors only needed for two-population analyses.

| **mu** | 4.6e^-9^ |
| --- | --- |
| **rho_over_theta** | 0.2 |
| **N_min** | 0 |
| **N_max** | 2,000,000 |
| **Tchanges_min*** | 0 |
| **Tchanges_max*** | 2,000,000 |
| **Tsplit_min**** | 0 |
| **Tsplit_max**** | 2,000,000 |
| **M_min**** | 0 |
| **M_max**** | 40 |

**Table S5**. Model choice for single-population inferences in DILS, for the three white-eye species and for the Grey-Headed Brown (GHB) form of the Reunion Grey White-eye. Columns headers indicate the different models contrasted, and the best model from random forest procedure is indicated in the first row (Note that the same models were chosen for the three species). Numbers indicate posterior probabilities of the best model.

|  | **constant *VS* contraction *VS* expansion** | **homo*N_e_* *VS* hetero*N_e_*** |
| --- | --- | --- |
|  | expansion | heteroN_e_ |
| Reunion Grey White-eye | 0.99 | 0.99 |
| *GHB form* | 1.00 | 1.00 |
| Mauritius Grey White-eye | 0.96 | 1.00 |
| Reunion Olive White-eye | 0.95 | 1.00 |

**Table S6**. Parameter estimation for single-population demographic scenarios in DILS, for the three white-eye species and for the Grey-Headed Brown (GHB) form of the Reunion Grey White-eye, with confidence interval (2.5%-97.5%). Three parameters were estimated, namely the population size prior to the population size change event (N_ANC_), the population size after this event (N_POP_), and the time since this event in number of generations (T_dem_).

|  | **ABC model parameters** | | | | | | | | |
| --- | --- | --- | --- | --- | --- | --- | --- | --- | --- |
|  | N_POP_ | | | N_ANC_ | | | T_dem_ | | |
|  | HPD2.5% | median | HPD97.5% | HPD2.5% | median | HPD97.5% | HPD2.5% | median | HPD97.5% |
| Reunion Grey White-eye | 90,514 | **101,936** | 117,948 | 33,928 | **36,910** | 40,200 | 15,712 | **20,842** | 26,763 |
| *GHB form* | 206,841 | **227,445** | 257,238 | 51,950 | **59,378** | 65,030 | 79,318 | **95,660** | 111,943 |
| Mauritius Grey White-eye | 399,051 | **493,676** | 604,792 | 62,638 | **66,032** | 70,205 | 38,544 | **44,146** | 49,889 |
| Reunion Olive White-eye | 633,251 | **707,856** | 871,491 | 111,604 | **128,383** | 143,798 | 207,529 | **246,493** | 286,294 |

**Figure S1**. Principal Component Analysis (PCA) of *ca* 13 million of SNPs that were extracted from sequence data from the three focal species: the Reunion Grey White-eye (purple), the Mauritius Grey White-eye (orange) and the Reunion Olive White-eye (green). The first axis separates the Reunion and Mauritius Grey White-eyes from the Reunion Olive White-eye, while the second axis separates the Reunion and the Mauritius Grey White-eyes from one another, as expected from phylogenetic relationships.

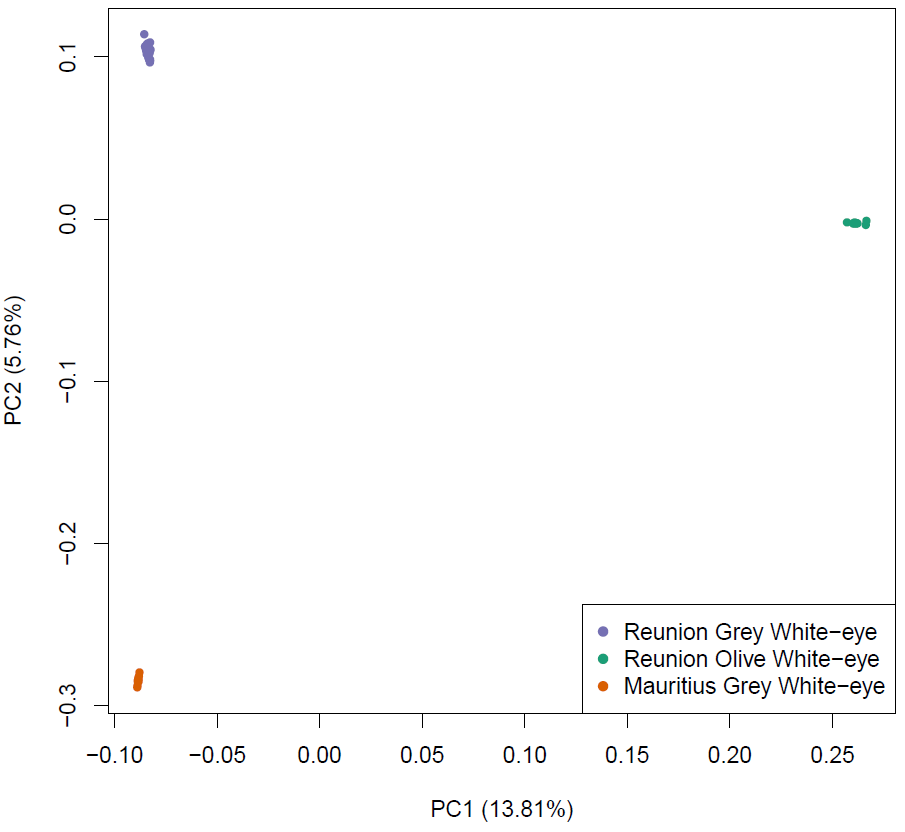

**Figure S2**. Schematic representations of: (A) the three single-population demographic models tested: population of constant size (constant), exponential potential decline at Tdem (bottleneck) and exponential growth at Tdem (expansion); (B) the four divergence models tested for two-population models in DILS: Strict Isolation (SI); Isolation with Migration (IM); Ancestral Migration (AM); Secondary Contact (SC). Red arrows indicate periods of gene flow. The models represented here are of constant size, but a demographic change can occur between present and T_split_ at a time T_dem1_ and T_dem 2_, changing the population sizes from N_ANC1_ to N_POP1_, and N_ANC2_ to N_POP2_ for both populations 1 and 2, respectively.

**A**
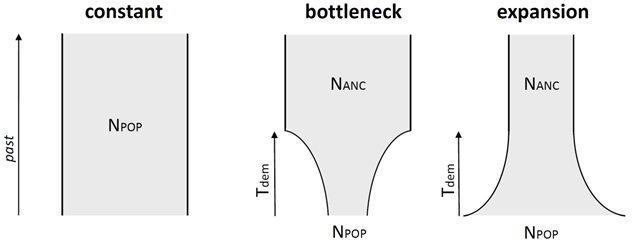

**B**
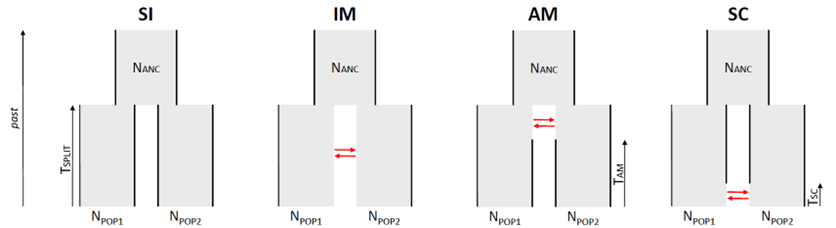

**Figure S3**. PSMC inferences of population size histories over time in the Reunion Grey White-eye. Colours indicate the different geographic forms, including the three lowland forms, namely the Grey-Headed Brown (GHB) form, the Brown-Naped Brown (BNB) form and the Lowland Brown-Headed Brown (LBHB) form, and the Highland form (HIGH). Vertical dotted lines and grey vertical bar indicate (from left to right): the Last Glacial Maximum 21,000 ya and a catastrophic eruption period in Reunion around 200,000 ya. Please note that the plots are in log-scales.

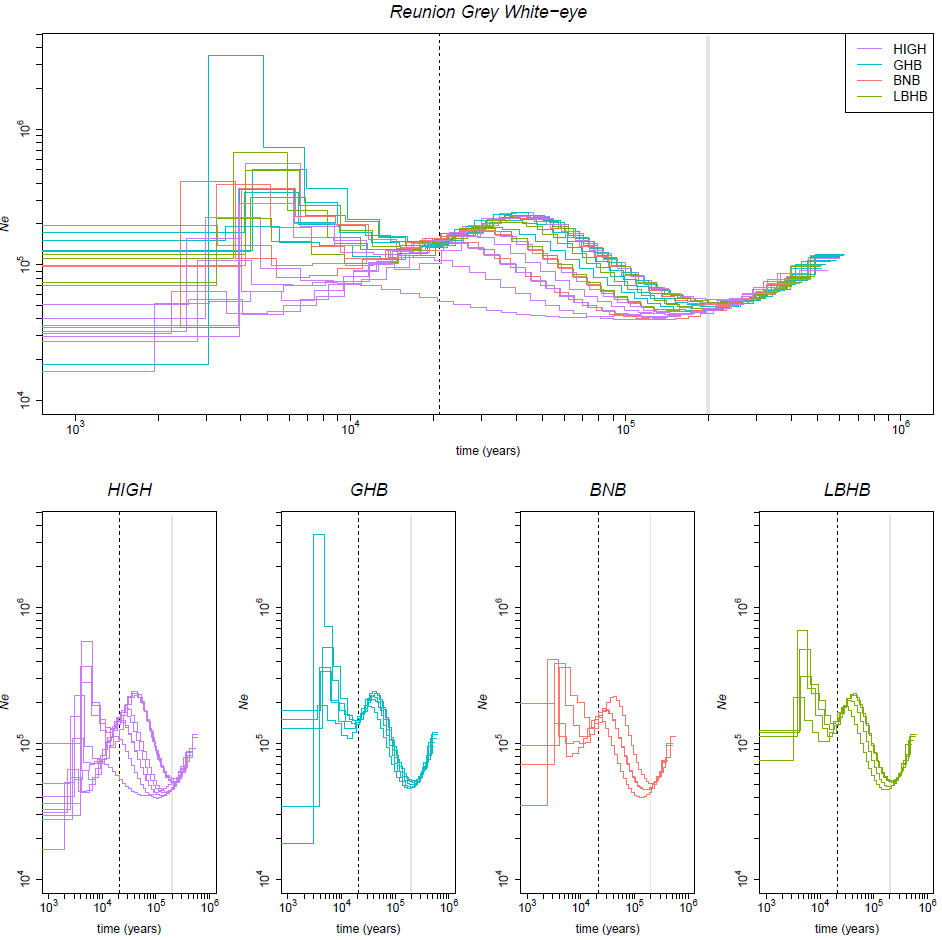

**Figure S4**. PSMC inferences of population size histories over time, varying the generation time (top line of the panel) from 1 year (middle) to 0.5 year (left) and 2 years (right), with a fixed mutation rate of 4.6e^-9^ mutations/site/generation and varying the mutation rate (bottom line of the panel) from 4.6e^-9^ mutations/site/generation (middle) to 3.4e^-9^ mutations/site/generation (left) and 5.9e^-9^ mutations/site/generation (right), with a fixed generation time of 1 year. See the section c of the methods for explanations on the choice of these values.

Colours indicate the three different species. Vertical dotted lines and grey vertical bar indicate (from left to right): the Last Glacial Maximum 21,000 ya and a catastrophic eruption period in Reunion around 200,000 ya. In Mauritius, the volcanic activity has been quite intense throughout the last 500,000 years but was never explosive during that period. Please note that the plots are in log-scales.

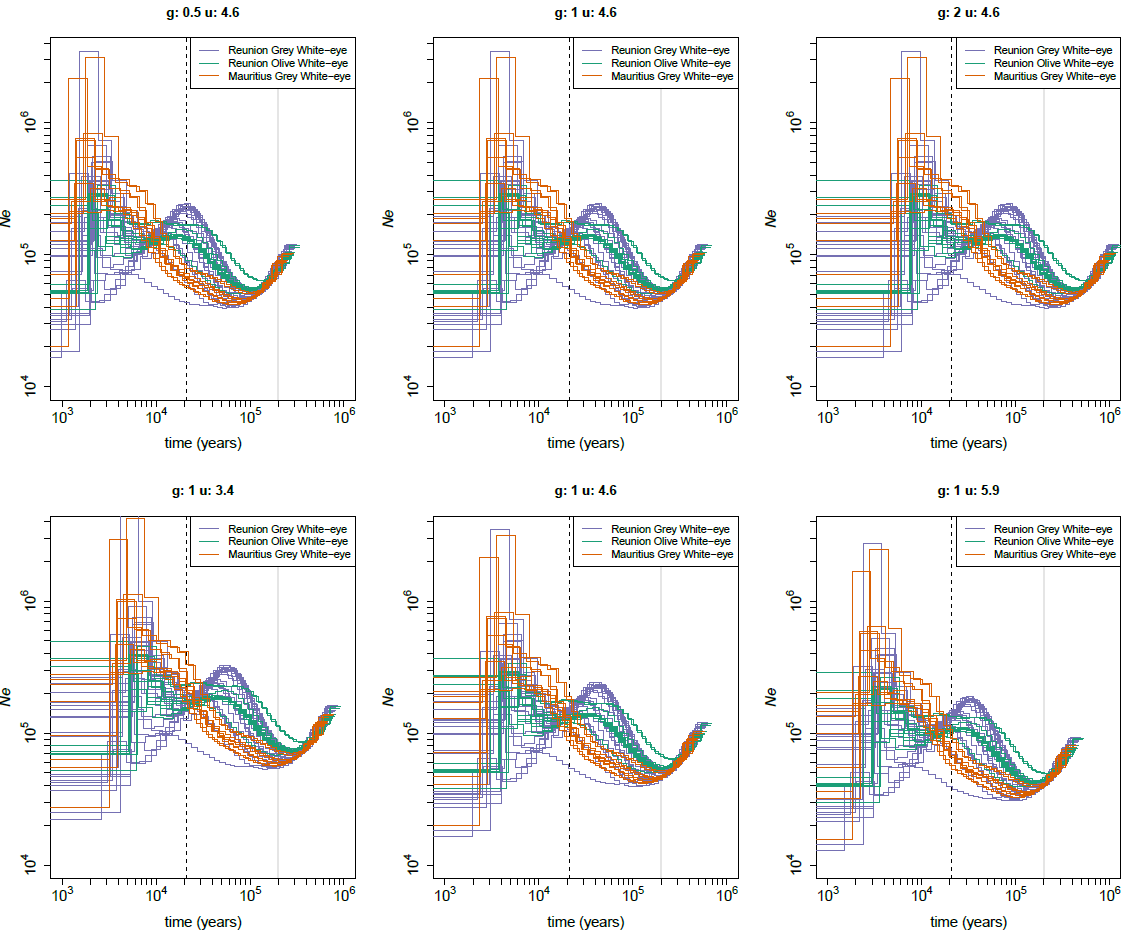

**Figure S5**. PSMC inferences of population size histories over time, using sequences with a minimal length of 1,000 pb (left) or 100,000 pb (right).

Colours indicate the three different species. Vertical dotted lines and grey vertical bar indicate (from left to right): the Last Glacial Maximum 21,000 ya and a catastrophic eruption period in Reunion around 200,000 ya. In Mauritius, the volcanic activity has been quite intense throughout the last 500,000 years but was never explosive during that period. Please note that the plots are in log-scales.

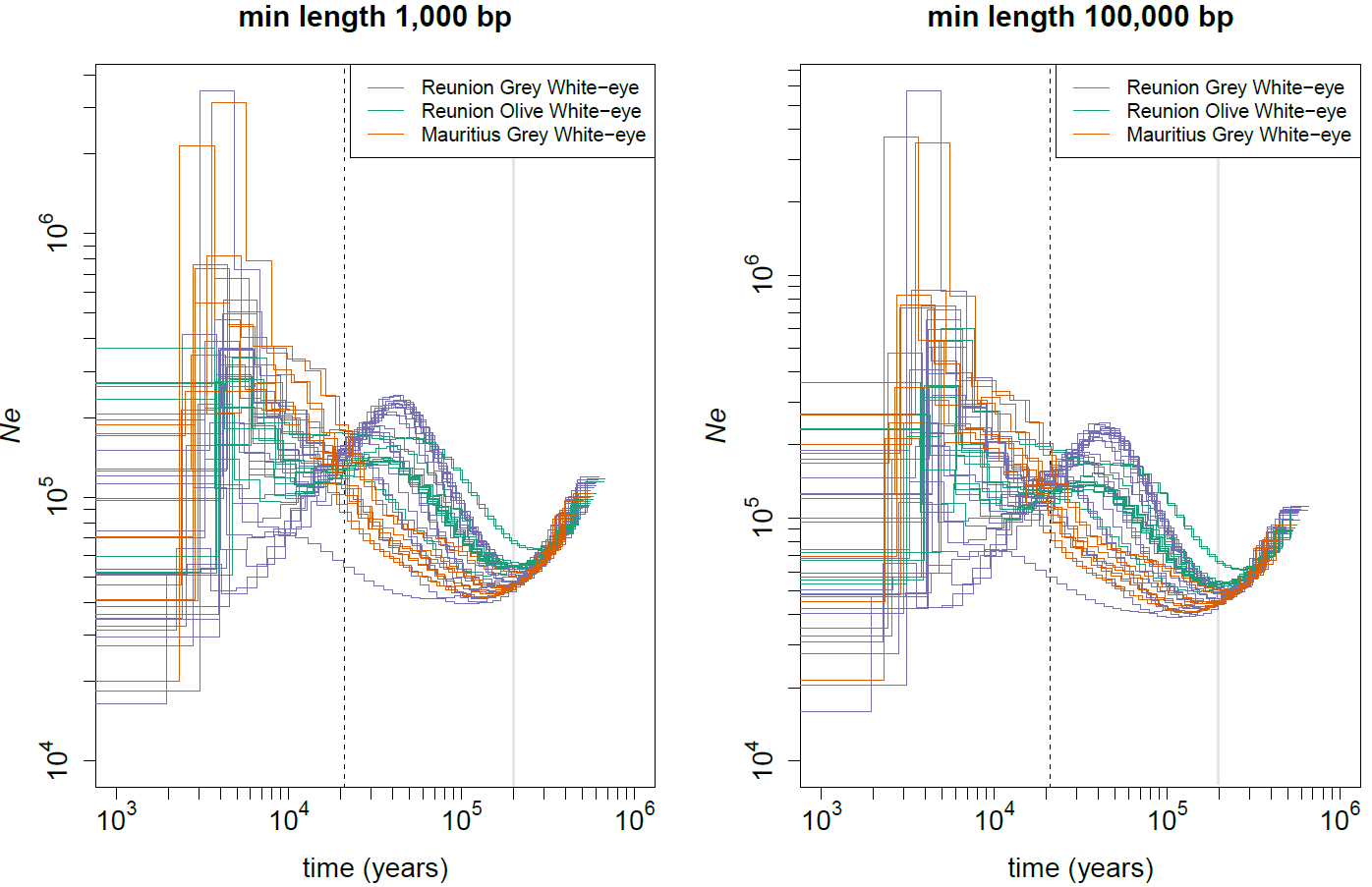

**Figure S6**. Schematic view of the DILS pipeline for two-population models. The three steps of model choices are depicted, with the best model in bold, and the number indicate the posterior probability of the selected model for the three pairs of species, from top to bottom: Reunion Grey White-eye – Mauritius Grey White-eye; Reunion Grey White-eye – Reunion Olive White-eye; Mauritius Grey White-eye – Reunion Olive White-eye.

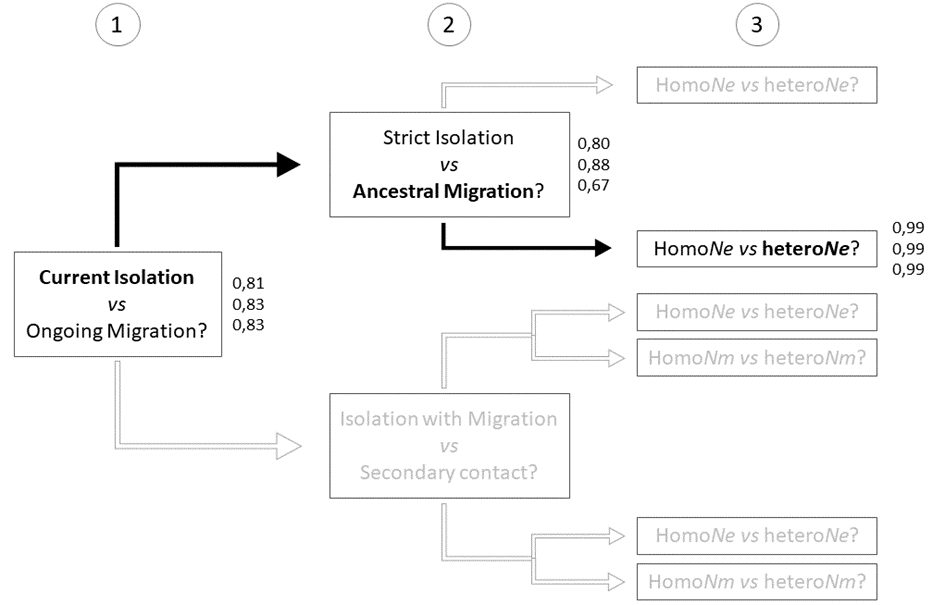
